# The renal response to FGF23 shifts from phosphaturia towards inflammation in kidney disease

**DOI:** 10.1101/2025.03.01.640954

**Authors:** Matthias B. Moor, Mikhail Burmakin, Anna Levin, Gül Gizem Korkut, David Brodin, Annika Wernerson, Annette Bruchfeld, Peter Bárány, Anna Witasp, Jaakko Patrakka, Hannes Olauson

## Abstract

**Background:** FGF23 excess is associated with morbidity and mortality, but the role of excessive circulating FGF23 concentrations as a mere biomarker or causative factor of pathology is controversial. Here, we investigated the consequences of FGF23 excess in kidney disease.

**Methods:** This study used three independent disease models: i) anti-glomerular basement membrane (Anti-GBM) disease in male C57BL/6 mice, ii) Adriamycin (doxorubicin)-induced nephropathy in female BALB/c mice, and iii) male DBA/2J mice fed an adenine-containing diet. Anti-GBM and Adriamycin mice and matched control mice received intravenous injections of recombinant FGF23 1µg or vehicle for six consecutive days (Anti-GBM) or once (Adriamycin model), with dissection 24h after the last injection. Adenine mice underwent organ harvesting after 15 weeks to establish ex vivo precision-cut kidney slices (PCKS) and 24h treatment with recombinant FGF23 or vehicle. In addition to histological and biochemical profiling, we assessed serum cytokines, biochemistry and renal transcriptomes and histology of mice and patients with IgA nephropathy. RNAseq data and published transcriptomes underwent gene set enrichment, bulk ligand-receptor interaction analysis and cell-type decomposition.

**Results:** Mice with Anti-GBM disease showed decreased glomerular filtration rate, albuminuria and renal tubular casts. FGF23 treatment increased phosphaturia, but also circulating soluble TNF receptor-1. Renal transcriptomes revealed FGF23-driven proinflammatory transcriptional signatures in murine Anti-GBM and also adverse Vcam1, Pdgfrb and chemokine ligand-receptor signaling in Anti-GBM but not in healthy mice. FGF23 increased transcriptome-inferred renal macrophage content in Anti-GBM mice. Findings were confirmed by immunofluorescence. In Adriamycin-induced nephropathy and in PCKS from the adenine nephropathy model, a short-term FGF23 excess caused expression of proinflammatory transcripts. Finally, human data revealed associations between histopathological or transcriptome-inferred renal immune cell infiltration and circulating FGF23 concentrations.

**Conclusion:** FGF23-driven patterns of proinflammatory gene and protein expression or leukocyte overabundance in the kidney were observed in several different models or states of FGF23 excess. The present data provide evidence that excess FGF23 directly drives inflammation in kidney disease and may serve as a therapeutic target.

## Introduction

Patients with chronic kidney disease (CKD) experience substantial morbidity due to earlier and more frequently occurring cardiovascular events compared to the general population, in part driven by uremic toxins, a systemic inflammatory state, endothelial dysfunction and vascular calcification. Vascular and other soft-tissue calcification is in part caused by the failure of the kidney to sufficiently excrete phosphate due to an end-organ resistance of the diseased kidney to the bone hormone fibroblast growth factor (FGF) 23.^1^ Circulating concentrations of FGF23 can rise up to 1000-fold in CKD, causing potential off-target effects and alterations in cellular signaling.^2^ Experimental studies in mice showed that FGF23 excess drives left heart hypertrophy via FGF receptor (FGFR) 4, calcineurin and nuclear factor of activated T cells (NFAT).^3–5^ Intriguingly, adverse effects of FGF23 excess on the heart are independent from systemic phosphate load, and FGF23 act synergistically with phosphate toxicity.^6,7^ In addition, FGF23 also causes hepatocytes to produce acute-phase proteins in calcineurin-dependent manner that may drive proinflammatory actions in other organs.^8^ However, renal adverse effects of FGF23 in CKD are less understood and, if they exist, underlie other mechanisms: Absence of FGFR4, the key mediator of cardiac adverse effects of FGF23, did not alter experimental CKD progression.^9^ Further, studies on adverse renal effects of FGF23 in mice revealed conflicting results: We and others previously reported that systemic *Fgf23* overexpression decreased renal expression of *Klotho*, a key co-receptor for FGF23 required for normal renal function.^10,11^ Exogenic delivery of FGF23 also induced kidney fibrosis and profibrotic cellular signaling in injury-primed renal fibroblasts.^11,12^ In contrast, completely binding FGF23 by antibody in rats increased CKD mortality.^13^ Further, *Fgf23* gene delivery had protective effects on endothelium and reduced AKI severity in mice with renal ischemia-reperfusion injury and contralateral nephrectomy.^14^

Among human studies, most observational data showed associations between circulating FGF23 and adverse cardiovascular outcomes, mortality and also adverse renal outcomes in several consortia.^3,15–17^. However, when assessing a potential causal effect by mendelian randomization among the general population, genetically predicted circulating FGF23 concentrations were not associated with renal function.^18^ Even more, a meta-analysis concluded that FGF23 is likely not causative for cardiovascular complications because of an absent exposure-response relationship, and because non-cardiovascular complications are also associated with FGF23 concentrations similar to cardiovascular complications.^15^

In the light of these controversies, it remains unresolved in the field if FGF23 is merely a biomarker or a real causative agent of morbidity such as CKD progression. With the intent of reducing this gap in knowledge, we performed experimental and observational studies on the adverse renal effects of FGF23 excess with a focus on CKD pathophysiology. Based on experimental glomerular disease with proteinuria-induced tubular injury, data from genetically induced FGF23 excess in mice, and biochemical and advanced transcriptome analyses in CKD patients, we here report that FGF23 induces proinflammatory cytokine expression and leukocyte migration to the kidney tissue in kidney disease. Further, in isolated precision-cut kidney slices we discovered kidney-intrinsic cytokine expression in response to FGF23 in murine CKD, thus providing evidence of liver-independent pro-inflammatory effects of FGF23 in kidney disease.

## Methods

### Animal experiments

Mice of C57BL/6J, DBA/2J and BALB/c strains were purchased from Charles River (France and Germany) or Scanbur (Denmark). Mice were housed in single-ventilated cages with 12 h light–12 h dark cycle and had *ad libitum* access to water and standard chow (Lantmannen, Kimstad, Sweden) unless indicated otherwise. Temperature was constant at 20±2 °C, and relative humidity was 50±5%. In all mouse experiments, euthanasia was performed by terminal exsanguination and cervical dislocation under anesthesia with isoflurane USP (Baxter), followed by organ collection.

### Kidney disease models

Within this study, we performed animal experiments using three mouse models of kidney disease:

1. Anti-glomerular basement membrane (Anti-GBM) disease was induced in C57BL/6 mice as described earlier.^19^ In brief, male mice aged 8 weeks were subcutaneously immunized with 400μl/kg of sheep IgG Freund’s complete adjuvant (F5881, Sigma-Aldrich). After four days, the mice were intravenously injected with 130 μl of nephrotoxic serum (NTS) obtained from Probetex (PTX-001S). Control mice were treated with vehicle (PBS in corresponding volume).
2. For induction of an Adriamycin nephropathy resembling human focal-segmental glomerulosclerosis (FSGS), we chose to extend investigation to female sex: Female BALB/c mice aged 8 weeks received a single intravenous injection of 10.5 mg/kg body weight Adriamycin (Sigma) that was diluted to 2 mg/mL in 0.9% NaCl2, or vehicle (0.9% NaCl2 in corresponding volume), derived from a protocol established earlier in the laboratory.^20^
3. A model of toxic tubulointerstitial kidney disease was established in male DBA/2J mice using adenine as described earlier.^21^ In brief, male DBA/2J mice aged 8 weeks were fed a diet on casein base containing 0.2% of adenine (SSNIFF Spezialdiäten GmbH, Germany). Throughout experiments, weight and overall appearance of all mice was scored 3 times per week. Spot urines and serum were collected at several time points during the study.

### Hormone injections

Treatments with FGF23 hormone were applied using a modification from an earlier protocol.^22^ Recombinant human FGF23 was obtained from R&D Systems, Minneapolis, MN, USA distributed via ThermoFisher (Cat. #100-52). Mice of the Anti-GBM model and controls were randomly allocated to receive daily intravenous injections with 1μg of FGF23 suspended in PBS-BSA 0.1% (Sigma) or PBSA-BSA 0.1% vehicle starting 1 day after receiving NTS. Mice of the Adriamycin model and corresponding controls were randomized for a single intravenous injection of 1μg of FGF23 or PBS-BSA 0.1% vehicle. Mice of both disease models were euthanized 24h after the last injection with FGF23.

### Glomerular filtration rate (GFR) assessment

To assess GFR in the Anti-GBM model at baseline and after 7 days after disease induction, a transdermal fluorescence detector (MediBeacon) was placed on the shaved back and was used to measure and quantify the clearance of injected FITC-sinistrin as previously described.^23^

### Precision-cut kidney slices (PCKS)

PCKS were established using a protocol modified from a previous study of the laboratory.^24^ In brief, a Compresstome® VF 310-0Z Vibrating Microtome (Precisionary Instruments, MA, USA) filled with Williams Medium E glutamax-I buffer (WME-solution Thermo Fisher, Waltham, Massachusetts, USA) containing 25 mM D-glucose (Merck, Darmstadt, Germany) was used to obtain 300µm kidney slices containing ca. 2/3 cortex and 1/3 medulla. The slices were transferred to 12 well plates and put to 1.5 ml of WME-solution at 37°C and 95% oxygen for 20 hours. Treatments with PBS-BSA 0.1% vehicle or recombinant human FGF23 (R&D Systems, #100-52) 200ng/mL, followed by harvesting and freeing in 1mL of RNAlater solution (Sigma Aldrich, Saint Louis, Missouri, USA), direct storage at −80°C, or fixing overnight in tubes containing 4% formaldehyde (Sigma Aldrich, Saint Louis, Missouri, USA).

### Serum and urine biochemistry assessment

Biochemical parameters were analyzed using quantification kits according to the manufacturer’s instructions, including blood urea nitrogen (BUN) by BUN Colorimetric Detection Kit (EIABUN, Thermo Fisher), creatinine by colorimetric detection using a kit (K625, Bio Vision) and phosphate by a colorimetric kit (Cat. MAK-030, Merk). Murine FGF23 was detected using ELISA kits for intact FGF23 (Immutopics, Inc., Cat. #60-6800) and C-terminal FGF23 (Immutopics, Inc., Cat. #60-6300), and human FGF23 was measured using an ELISA kit for intact FGF23 (Immutopics, Inc., Cat. #60-6600) as per the manufacturer’s instructions.

### Immune protein array

For quantification of 40 inflammation-related proteins in mouse serum, two Mouse Inflammation Array C1 kits (RayBiotech, #AAM-INF-1-8) were used as per the manufacturer’s instructions, with 60µL serum from up to three mice pooled per sample, for a total of 4 experimental groups with 4 independent biological replicates per group. Membranes were bathed in the kits’ chemoluminescence reagents for 30 seconds. Images of the membranes were obtained on an Odyssey® XF Imaging System (LICORbio). Spots on digital images underwent densitometric quantitation using an ImageJ Protein Array Analyzer macro,^25^ with standard settings and linear background subtraction. The obtained data were analyzed using R/limma with Benjamini-Hochberg correction for multiple testing and interaction analysis in two-factorial design (factors: treatment with FGF23 vs. vehicle, and disease state: Anti-GBM vs. healthy).

### Histology

Organs were fixed in 4% formaldehyde and underwent paraffin embedding. Four µm thick sections were made, followed by standard staining using Hematoxylin & Eosin, or Masson’s Trichrome staining kit (Cat. #HT15, Sigma-Aldrich). Brightfield images were scanned using a ZEISS Axioscan 7 scanner. The presence or absence of renal tubular protein casts was determined by the following score: 0, no casts; 1, >0-20%; 2, 20-40%, 3, 40-60%; 4, 60-80%; 5, >80% of the tubules are affected.

### Immunofluorescence

Paraffin-embedded sections were dewaxed and rehydrated, followed by antigen retrieval at 98°C for 20min in 10mM Tris, 1mM EDTA, 5% Tween. Subsequently, slides were washed with PBS before blocking with Dako serum-free protein block (X0909, Agilent) for an hour at room temperature. Followingly, slides were incubated overnight at 4°C with primary antibodies against CD45 (Abcam ab61100, 1:100) or F4/80 (Invitrogen Cat. #14-4801-82, 1:50). Alexa Fluor-conjugated secondary antibodies were from Invitrogen. Slides were co-stained with Hoechst 33342 (H3570, 1:10,000, Life Technologies) for 10_min at room temperature. Sudan black B (0.1% in 70% Ethanol, Cat. #199664, Sigma-Aldrich) was applied for 25_min at room temperature followed by mounting in Dako fluorescent medium (S3023, Agilent). Slides were scanned at 20X magnification using a Vectra 3 automated quantitative pathology imaging system (Akoya Biosciences). Subtraction of the background signals and image export were performed using inForm® software (Akoya Biosciences). Images were exported for quantification of stained cells using QuPath version 0.5. After automated cell detection, the fraction of CD45 positive immune cells underwent automated quantification using an intensity threshold, and F4/80-positive macrophages underwent manual identification by a blinded investigator, while comparing to the autofluorescence channel. For CD45 staining, the fraction of positive cells of approximately 7000 cells per animal were reported, and for F4/80 ca. 3’000 cells per animal.

### RNA isolation

A fragment of the renal pars intermedia per mouse was homogenized and lyzed using a TissueLyser (Qiagen) in 600 µl RLT buffer + DTT. RNA was then extracted using Qiagen RNeasy mini kit (Qiagen) according to manufacturer’s instructions.

### RNA-seq (Illumina mRNA ligation kit)

Total RNA was subjected to quality control with Agilent Tapestation according to the manufacturer’s instructions. To construct libraries suitable for Illumina sequencing the Illumina TruSeq Stranded mRNA ligation. A sample preparation protocol including cDNA synthesis, ligation of adapters and amplification of indexed libraries was used. The yield and quality of the amplified libraries was analyzed using Qubit by Thermo Fisher and the Agilent Tapestation. The indexed cDNA libraries were normalized and combined. The pools were sequenced on the Illumina Nextseq 2000 with a P3 100 sequencing run generating 58 bp paired-end reads.

### Bulk RNAseq data analysis

Demultiplexing was performed using bcl2fastq (v2.20.0). Reads were aligned GRCm38 from Ensembl using STAR (v2.7.9a). Aligned reads were assigned to coding regions of various biotypes defined in Mus_musculus.GRCm38.102.gtf from Ensembl using featureCounts (v1.5.1). Annotations were collected from Ensembl BioMart using the biomaRt package. All comparisons were performed using the DESeq2 package. For Anti-GBM and Adriamycin experiments, the 3 contrasts disease status (kidney disease model vs. healthy controls), treatment status (FGF23 vs. vehicle), and the interaction between disease status and treatment status were tested, with false discovery rates adjusted for the 3 contrasts combined by the Benjamini-Hochberg method. Gene set enrichment analysis (GSEA) was performed using the fgsea package with the Molecular Signatures Database (MSigDB) collection of gene sets. Ligand-receptor interaction analysis was performed using BulkSignalR v.0.0.9^26^ using differential mode.

### Transcriptome dataset retrieval from public repositories

Publicly available microarray datasets of renal gene expression were retrieved from Gene Expression Omnibus Dataset Browser, including kidney dataset GDS3361 of 8-weeks old male *Fgf23* transgenic and control mice by Marsell et al.^27^ and kidney dataset GDS879 of 5-weeks old sex-balanced (60% males) carrying the Hyp mutation in the *Phex* gene.^28^ Single-cell RNAseq data were downloaded from Gene Expression Omnibus, including GSE107585 of murine kidney from 7 sex-mixed healthy C57BL/6 mice by Park et al.^29^

### Human cohort of patients with IgA nephropathy

We reanalyzed published data from 59 adult patients with IgA nephropathy (IgAN) from the Karolinska Kidney Biopsy Project in relation to demographic and clinical characteristics, medication data and serum biochemistry^30^. In addition, the diagnostic kidney biopsy evaluations were accessed, including Oxford classification, and we obtained the processed RNAseq count data from micro-dissected non-glomerular (tubular) fractions of the kidney. The RNAseq analysis procedure including filtering and normalization is described in Levin et al.^30^

### Bulk transcriptome deconvolution

For analyses of relative renal segment or cell type abundance, single-cell RNAseq dataset GSE107585 was reanalyzed using Seurat 5.0.3. After excluding cells with less than 200 or more than 2500 RNA counts or cells with >30% mitochondrial RNA, normalization and clustering, clusters were manually annotated based on cluster-specific expression and markers with reference to the original publications of the datasets.^29,31^ Subsequently, Bisque v.1.0.5^32^ was used in reference-based decomposition mode for deconvolution of bulk RNAseq or microarray datasets.

### Statistics

Data analyses and visualizations were made with RStudio 2023.12.1 and/or GraphPad Prism 8 or SuperPlotsOfData.^33^ Frequency data were analyzed by Chi-square test, and continuous data by t-test, paired t-test or Two-way ANOVA as appropriate. Univariable and multivariable linear regression models were used with dependent variables as indicated, using adjustments for immune-relevant confounders for multivariable models, namely 25OH-vitamin D, PTH, corticosteroid use (yes/no) and other immunosuppressants (yes/no). In the models, we also included an interaction term of measured GFR and cell abundance. Circulating immune proteins and RNAseq data were analyzed in 2-factorial design using limma and DESeq2, respectively, as reported above.

### Study approval

All animal experiments were carried out at Karolinska Institutet following the protocols approved by the Ethical Committee on Research Animal Care (Linköping Animal Experimentation Ethics Committee; DNR 1336). The Karolinska Kidney Biopsy Project was approved by the Swedish Ethical Review Authority. The reanalysis of existing transcriptomes in the public domain required no authorization.

### Data availability

The bulk RNAseq datasets generated within this study are accessible at Gene Expression Omnibus under the accession number GSE272289. Existing single-cell RNAseq data were retrieved from Gene Expression Omnibus by the accession number GSE107585. Existing renal microarray transcriptome datasets GDS3361 and GDS879 were retrieved from GEO Dataset Browser. Analyses and visualizations were made using adaptations of existing code from the tools <u>BulkSignalR</u>, <u>Seurat</u>, <u>scCustomize</u>, <u>bisque</u>, <u>Enhanced Volcano</u>, and <u>SuperPlotsOfData</u>. Data from the Karolinska Kidney Biopsy Cohort cannot be deposited to public repositories due to ethical restrictions but are available upon request; requests for access to the data can be put to our Research Data Office at Karolinska Institutet.

## Results

### Partial renal resistance of the phosphaturic response to FGF23 in mice with Anti-GBM disease

Previous work by us and others has shown systemic proinflammatory responses occurring under FGF23 excess in mice.^8,10^ In the present study, we aimed to substantiate the renal consequences of such adverse effects of FGF23 excess, using two mouse models with proteinuria-induced proximal renal tubular dysfunction. We first induced Anti-GBM disease in adjuvant-immunized C57BL/6 mice using nephrotoxic serum (**Figure 1A**) as previously described.^19^ Mice additionally received daily intravenous administrations of recombinant FGF23 or vehicle over 6 days (**Figure 1A**). We have previously assessed the morphological effects of this protocol of Anti-GBM disease induction using electron microscopy of the kidney, in which we detected severe podocyte effacement.^19^ In the present study, we first verified the validity of the model by the decreased GFR at day 7 (**Figure 1B**), increased plasma creatinine and BUN (**Supplemental Table 1**), the increased urinary albumin / creatinine ratio in the Anti-GBM groups (**Figure 1C**), and by the tubular protein precipitates (casts) on renal histology resulting from proteinuria (**Figure 1D**).

**Figure 1.**
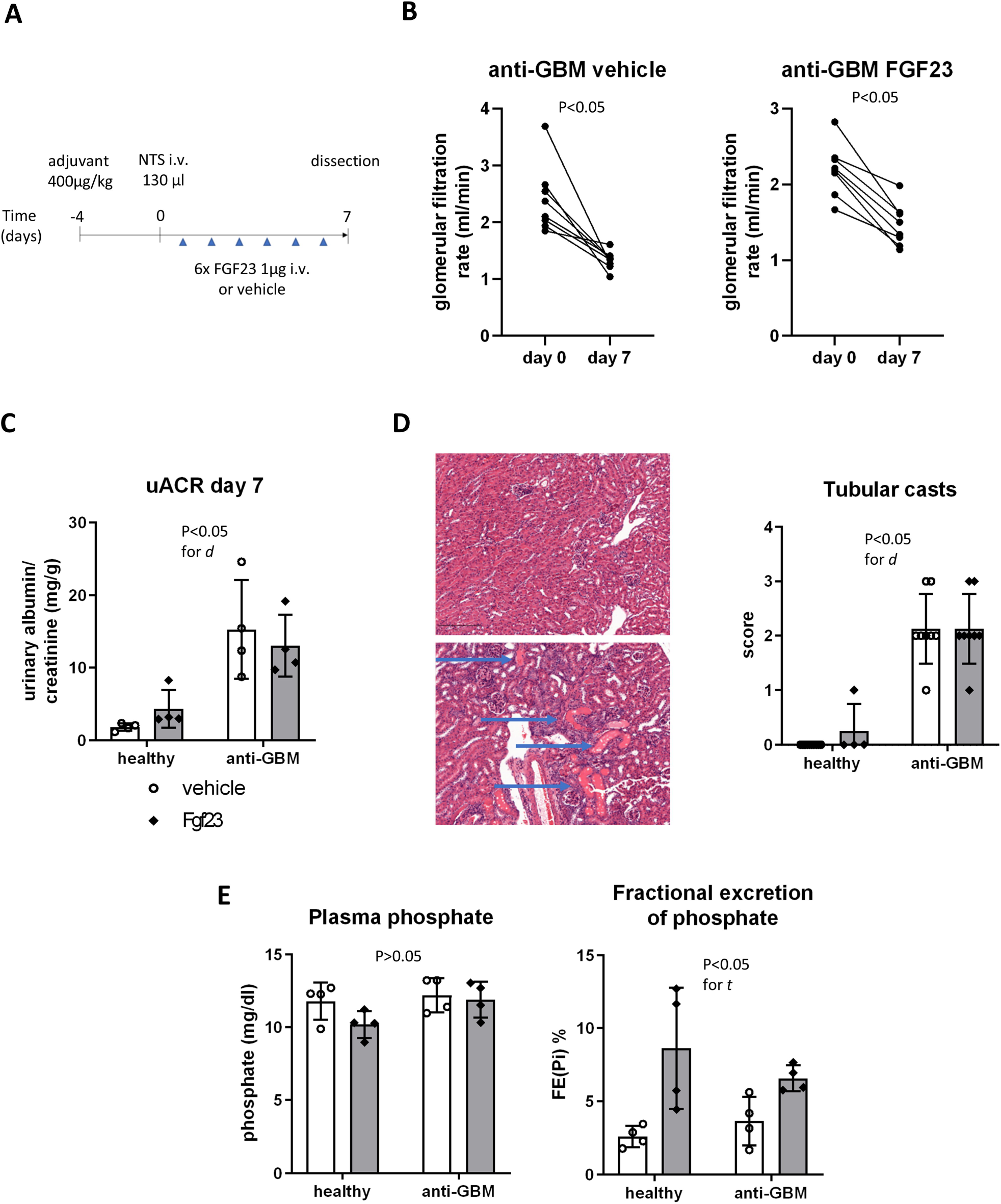
Anti-GBM disease causes tubular damage and partial renal resistance to FGF23. A depicts experimental workflow in male C57BL/6 mice undergoing induction of anti-glomerular basement membrane (Anti-GBM) disease using nephrotoxic serum (NTS) followed by 6 days of intravenous (i.v.) FGF23 or vehicle treatment. B show glomerular filtration rate (GFR) of different experimental groups at day 0 and 7. C shows urinary albumin/creatinine ratio (uACR) at day 7. D shows example renal sections negative or positive for renal tubular casts (arrows) and quantitative tubular cast scores, E plasma phosphate and fractional excretion of phosphate of health mice and mice with Anti-GBM treated with vehicle or FGF23. Analyses in B: paired t-test. C-E: Two-way ANOVA. d, disease state. t, treatment.

Next, we determined the physiological effects of the six administrations of recombinant FGF23 or vehicle. To this end, we determined the concentrations of circulating FGF23. FGF23 was elevated 24h after the last injection of both healthy and diseased FGF23 groups (**Supplemental Table 1**). Further, the FGF23 administrations tended to decrease plasma phosphate compared to vehicle in healthy mice (**Figure 1E**). The fractional excretion of phosphate (FE_Pi_) was increased by FGF23 compared to vehicle, with around three-fold increase of FE_Pi_ in healthy mice and a two-fold increase in Anti-GBM mice (**Figure 1E**). However, FGF23 did not directly further aggravate renal function in the Anti-GBM disease model, as GFR was comparable at day 7 between the Anti-GBM groups treated with FGF23 or vehicle (p>0.05, **Figure 1C-D**). Overall, the present data demonstrated substantial kidney disease in the Anti-GBM model, and evidence of a partial renal resistance to the phosphaturic actions of FGF23 in Anti-GBM disease.

### FGF23 excess triggers proinflammatory renal responses and increased immune cell abundance in the diseased kidney

FGF23 excess caused by recombinant hormone administration^22,34^ or by *Fgf23* overexpression^27^ induces renal cell signaling via FGF receptors 1 and 3 and mitogen-activated protein kinase (MAPK) phosphorylation leading to gene expression of *Cyp24a1* encoding a vitamin D metabolizing enzyme. In mice with glomerular disease, however, FGF23 induced renal injury marker *Lcn2*.^34^ To globally profile the transcriptional changes, we performed unbiased bulk RNAseq in healthy mice and the mice with Anti-GBM disease treated with FGF23 or vehicle for six days. Overall, we found no differentially expressed genes (DEG) following 24 hours since the last FGF23 injection in healthy mice, but 27 genes were upregulated, and 2 genes downregulated by FGF23 in Anti-GBM (**Figure 2A, Supplemental Table 2**). Of note, the expression of renal FGF23 co-receptor *Klotho* in the Anti-GBM groups remained at 70% compared to the healthy groups (**Supplemental Table 2**). In an interaction analysis, we next determined how FGF23 effects differ between Anti-GBM disease and healthy mice, revealing 23 DEGs (**Figure 2A, Supplemental Table 2**). **Figures 2B-C** show the transcripts with a difference in FGF23 effect between Anti-GBM and healthy mice, including mRNA of *Tgtp1* and *Tgtp2* encoding T-cell specific GTPases, and *Irf1* encoding an interferon regulatory factor. Next, we performed a gene set enrichment analysis (GSEA) to gain insight into the overall pattern of affected transcripts. GSEA revealed proinflammatory FGF23-dependent transcriptional signatures in Anti-GBM vs. healthy mice, including an enrichment of interferon signaling (**Figure 2D-F, Supplemental Figures 1A-C**).

**Figure 2.**
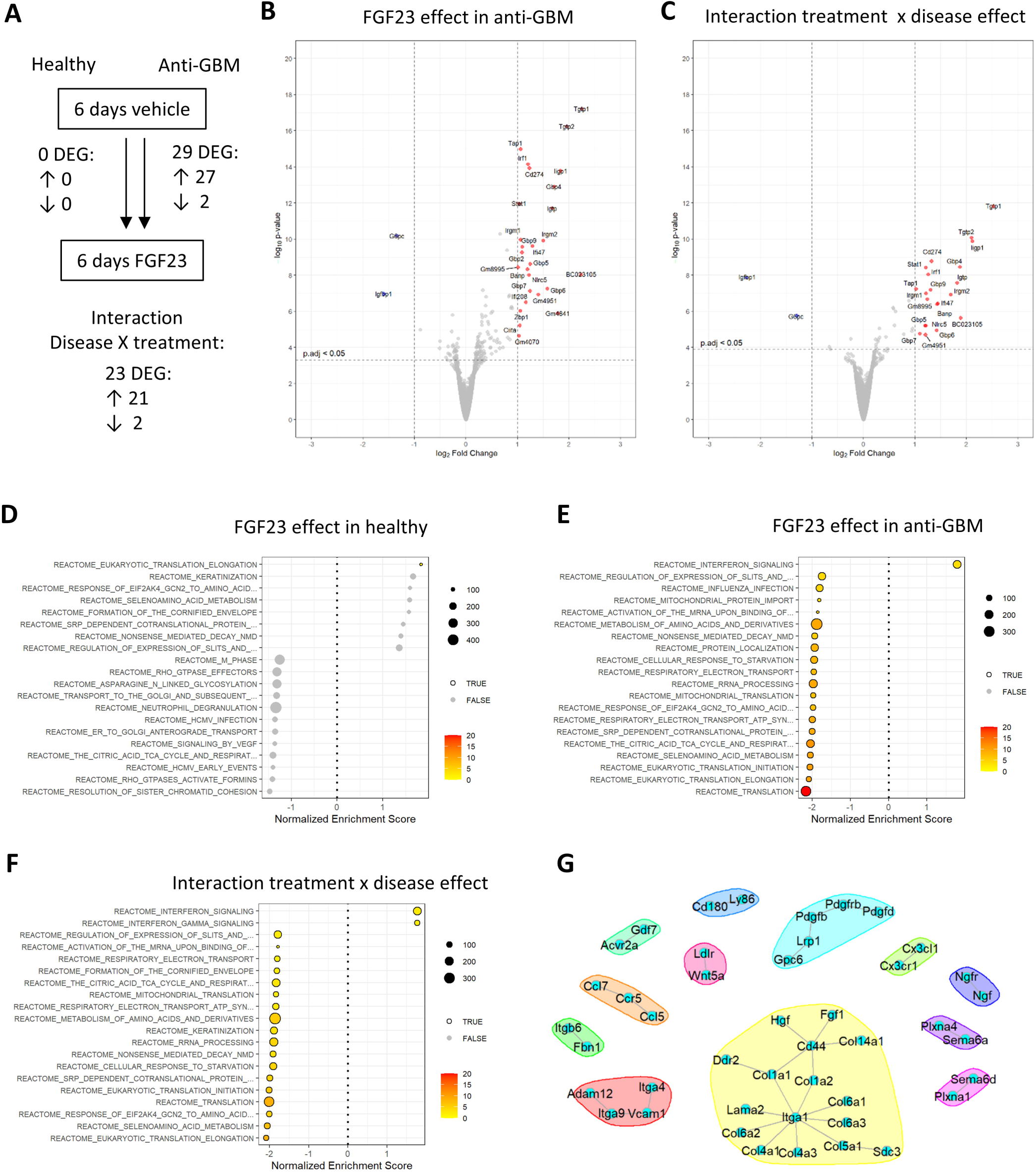
Six day course of FGF23 treatment induces renal transcriptional signatures of inflammatory responses and injury. A indicates the number of differentially expressed genes (DEG) according to experimental comparison in renal bulk RNAseq. B-C depict Volcano Plots of DEGs above a cut-off of adjusted p<0.05 and log2-fold change >1, in comparison of FGF23 vs vehicle effect in mice with Anti-GBM (B) and the interaction between treatment and disease effect (C). D-F depict significant Reactome gene set enrichment analyses of FGF23 effects in healthy mice (D), mice with Anti-GBM disease (E) and the interaction between treatment and disease effect (F). G depicts a network of ligand-receptor interaction pairs that were significant for FGF23 vs vehicle comparisons in mice with Anti-GBM disease by bulk RNAseq. Anti-GBM, anti-glomerular basement membrane disease. N=3 for Anti-GBM groups and n=4 for healthy groups.

Next, we aimed to better understand the mechanisms and involved signaling molecules by which FGF23 induced the observed proinflammatory renal transcriptional signatures. To this end, we performed ligand-receptor interaction analysis of the bulk RNAseq data. This analysis revealed that the FGF23 effect in the Anti-GBM model included on the one hand signaling by several chemokines of the classes CC, CXC and CX3C and their corresponding receptors (**Figure 2G, Supplemental Figure 2D**). On the other hand, the ligand-receptor interaction analysis provided evidence of FGF23-induced cell signaling by collagen-binding integrins, Toll-like receptor and TGF-beta signaling, and the involved molecules *Vcam1*, *Pdgfrb* and *Fbn* are all markers of injury and fibrogenesis (**Figure 2G, Supplemental Figures 2D-E**). In summary, in-depth bulk transcriptome analyses provided several layers of evidence pointing towards pro-inflammatory renal actions of FGF23 in the Anti-GBM disease model.

Based on the above findings, we next wondered to which extent FGF23 may affect proinflammatory cytokines at the protein level. Therefore, we profiled 40 circulating immune proteins by a protein array in sera of healthy and Anti-GBM mice treated with vehicle or FGF23 (**Supplemental Figure 3**). In healthy mice, FGF23 increased the circulating concentrations of soluble tumor necrosis factor (TNF) receptor (sTNFR) 1 (**Figure 3A**). By contrast, in mice with Anti-GBM disease, we found no FGF23-induced alterations in immune proteins (**Figure 3B**). Consequently, a two-factorial analysis of the interaction between treatment and disease state revealed sTNFR1 as the sole significantly altered immune protein (**Figure 3C**). Further, when comparing all animals regardless of FGF23 treatments, sTNFR2 was increased in Anti-GBM compared to healthy mice (**Figure 3D**). The individual results of sTNFR1 and sTNFR2 in each experimental group are shown in **Figure 3E-F**, and all comparisons in **Supplemental Table 3**. Overall, Anti-GBM disease in mice already induced a proinflammatory milieu in serum with elevated sTNFR1 and sTNFR2 concentrations similarly as in human CKD.^35,36^ Consequently, FGF23 had an effect on sTNFR1 only in disease-free mice but did not alter the already elevated circulating sTNFR1 in mice with Anti-GBM.

**Figure 3.**
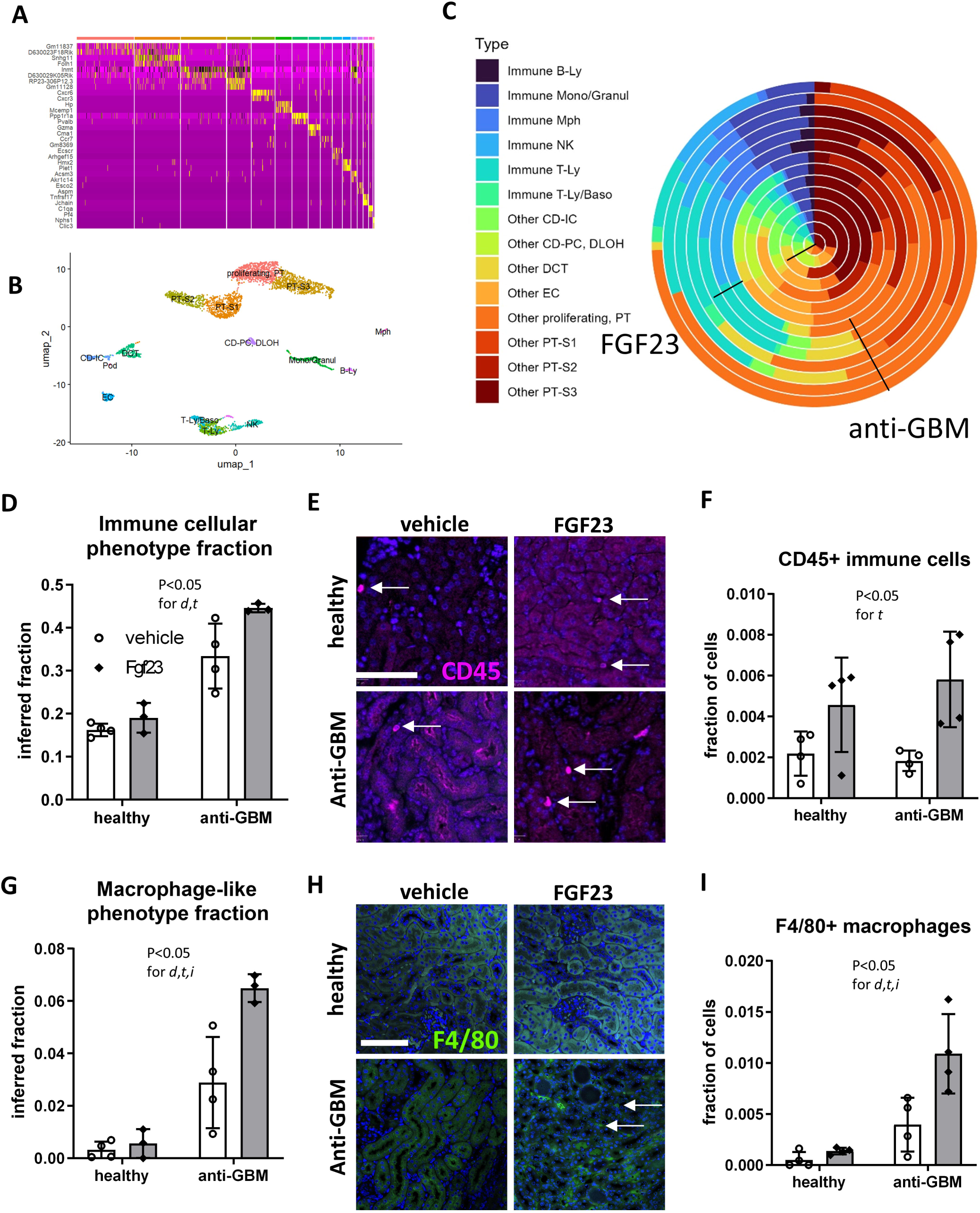
Bulk RNAseq deconvolution and immunofluorescence staining reveal FGF23-driven increase in overall immune cell and macrophage abundance in kidney of mice with Anti-GBM disease. A-B depict the annotation of renal cell clusters in reanalysis of single-cell RNAseq dataset GSE107585 of murine kidney from 7 sex-mixed healthy C57BL/6 mice, see also Supplementary Figure 2. C shows a wedding pie plot of the bulk deconvolution of the renal cellular composition according to FGF23 treatment and Anti-GBM disease state as indicated by labels. Renal overall immune cells and macrophage-like cells are displayed by bulk deconvolution (D, G) and by immunofluorescence with automated quantification for CD45 (E-F) and F4/80 (H-I). RNAseq: N=3 for Anti-GBM groups and n=4 for healthy groups. Immunofluorescence: n=4 per group. Statistical analysis: Two-way ANOVA. d, disease state. t, treatment. i, interaction. Anti-GBM, anti-glomerular basement membrane disease. Ly, lymphocyte. Mono, monocyte. Granul, granulocyte. Mph, macrophage. NK, natural killer cell. Baso, basophil. CD, collecting duct. IC, intercalated cells. PC, principal cells. DLOH, descending limb of Henle. DCT, distal convoluted tubule. EC, endothelial cell. PT, proximal tubule. S, segment.

In view of the renal transcriptional signatures and cytokines suggesting that FGF23 promotes a proinflammatory environment, we aimed to determine if FGF23 caused an influx of immune cells to the kidney. Thus, we determined the relative abundance of different renal cell compartments by bulk transcriptome deconvolution and by immunofluorescence (**Figure 4**). For bulk deconvolution, we first reanalyzed and annotated existing single-cell RNAseq data of kidney from healthy C57BL/6 mice (**Figure 4A, Supplemental Figure 4**). Second, we inferred the fraction of major cell types, including total immune cells and specifically macrophages from renal bulk transcriptomes of healthy and Anti-GBM mice treated with vehicle or FGF23 for 6 days (**Figure 4C**). FGF23 excess provided an additional increase in total immune cells (**Figure 4D**) and specifically macrophages (**Figure 4D**) in addition to an observed increase in immune cells induced by Anti-GBM disease. Next, we validated these transcriptome-derived findings using immunofluorescence staining of the kidney, confirming an increase in overall CD45-positive immune cells (**Figure 4E-F**) and F4/80-positive macrophages (**Figure 4G-H**) in FGF23-treated mice compared to vehicle-treated mice with Anti-GBM disease. Together, these findings provide strong evidence of an FGF23-driven increase in immune cell content in the diseased kidney.

**Figure 4.**
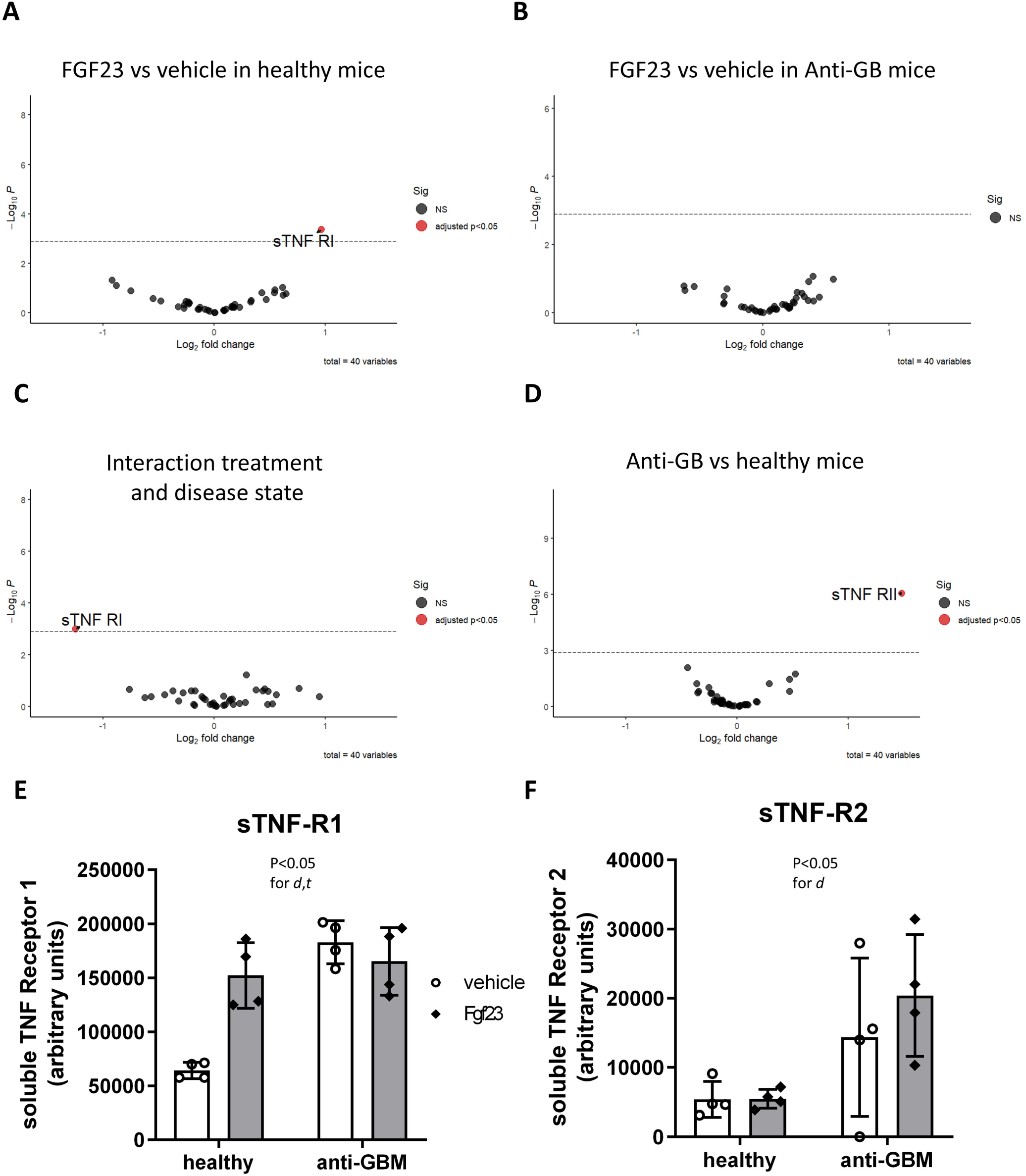
Immune protein profiling highlights an increase in circulating soluble TNF Receptors induced by FGF23 and Anti-GBM disease in mice. A plasma cytokine protein array shows FGF23 effects in healthy male C57BL/6 (A) and mice treated with nephrotoxic serum to induce Anti-glomerular basement membrane (Anti-GBM) disease (B). Interaction between treatment and disease state (C) and overall disease effect (D) are shown. E-F depict analyses of soluble TNF Receptors (sTNF-R) 1 and 2 by Two-way ANOVA. d, disease state. t, treatment. N=4 biologically independent replicates per group.

### Time-dependent and renal-intrinsic proinflammatory effects of FGF23

With evidence of excessive FGF23 causing proinflammatory effects in the diseased kidney, we aimed to determine the time dependency of this aberrant FGF23 effect. First, we used bulk transcriptome deconvolution of existing microarray data from kidneys of mice transgenically overexpressing Fgf23 and mice harboring a mutation in the *Fgf23* repressor gene *Phex* (Hyp mouse) and their respective controls, as models of long-term FGF23 excess. This analysis revealed that genetically induced *Fgf23* excess tends to increase the total renal immune content (**Figure 5A**) and significantly increases the inferred renal macrophage content (**Figure 5B**). Conversely, we also studied mice with a short-term treatment of FGF23 in a different disease model of Adriamycin nephropathy (**Figure 5C**): We first induced glomerular and renal tubular disease using Adriamycin in female mice, leading to an over 8-fold increase in urinary albumin/creatinine ratio (ACR) (**Figure 5D**). In addition, serum BUN tended to be higher and serum creatinine was significantly increased in mice with Adriamycin-induced nephropathy (**Supplemental Table 1**). We then administered a single intravenous injection of recombinant FGF23 24h prior to euthanasia in both healthy mice and mice with Adriamycin-induced nephropathy. There was no change in circulating FGF23, phosphatemia or urinary phosphate excretion at the 24h interval (**Supplemental Table 1**). In renal RNAseq analysis of these mice (**Figure 5F-H**, **Supplemental Table 4**), the top DEGs in an interaction analysis between FGF23 treatment and disease state included proinflammatory transcription factor, *Cebpb,* and an increased DNA damage transcript, *Dclre1b* (**Figure 5H**, **Supplemental Table 4**). GSEA pointed towards FGF23 decreasing chaperone-mediated autophagy FGF23 in diseased kidney, with substantially fewer overall effects induced by a single FGF23 injection at the 24h interval (**Figure 5F-G**). Overall, a long-term genetic induction of FGF23 excess increased renal macrophage content in otherwise healthy mice, whereas a short-term FGF23 treatment of 24h induced only limited potential proinflammatory FGF23 effects in mice with kidney disease and none in healthy mice.

**Figure 5.**
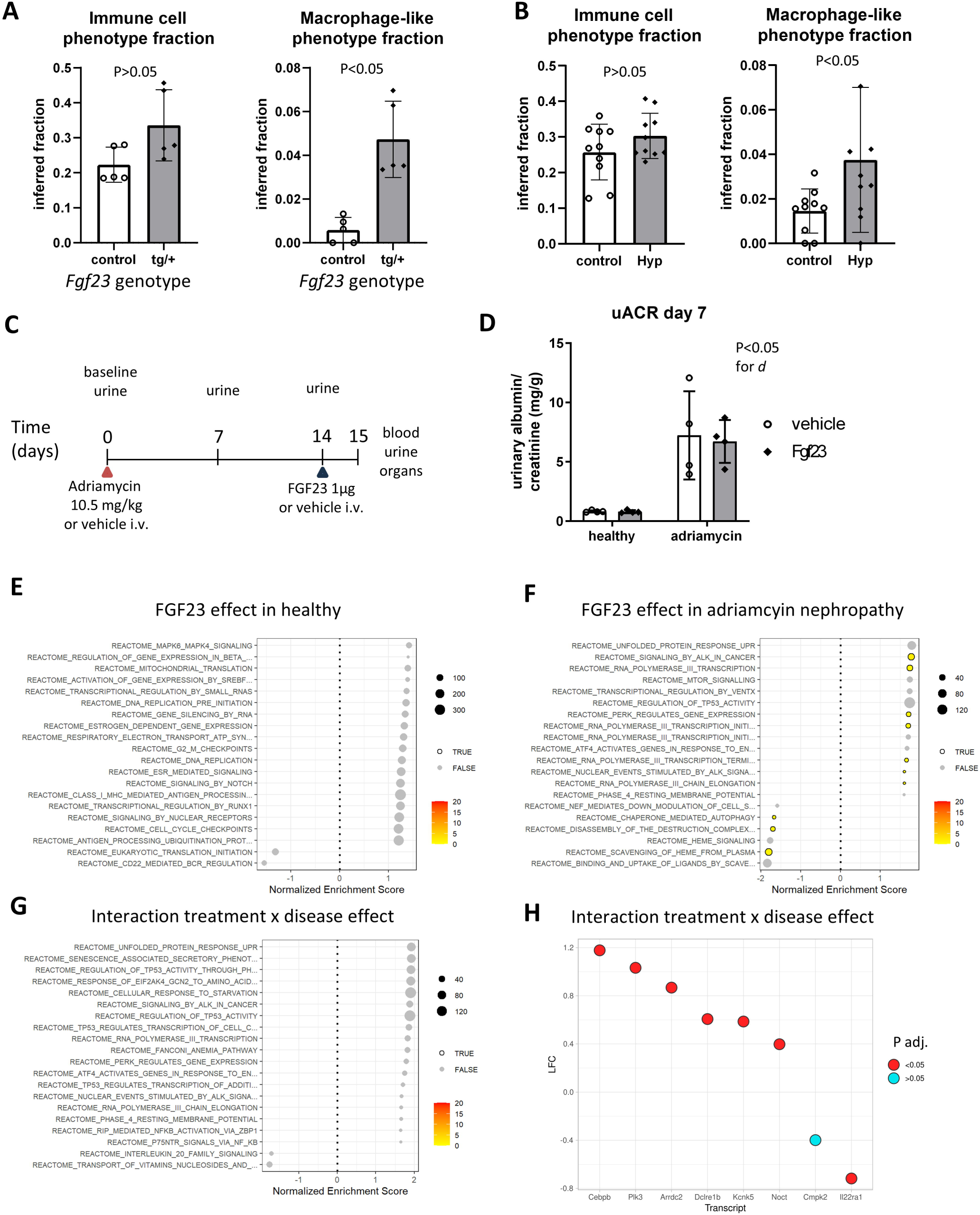
Renal immune cell recruitment driven by FGF23 excess is exposure time-dependent. Renal microarray transcriptome datasets GDS3361 of male *Fgf23* transgenic and control mice or sex-matched Hyp mice of dataset GDS879 (B) underwent bulk deconvolution with reference to single-cell RNAseq dataset GSE107585 of murine kidney from 7 sex-mixed healthy C57BL/6 mice and, followingly, visualization of overall fractions of inferred immune cells and macrophage-like cells as indicated. C shows the experimental workflow of experiments with female BALB/c mice undergoing Adriamycin (doxorubicin) nephropathy followed by a single intravenous (i.v.) injection of FGF23 or vehicle. D shows urinary albumin/creatinine ratio (uACR) 7 days after induction of Adriamycin nephropathy. Statistical analysis: Two-way ANOVA. d, disease state. E-G show significant Reactome gene set enrichment analyses of renal FGF23 effects in healthy mice (E), mice with Adriamycin nephropathy (F) and the interaction between treatment and disease effect (G). H depicts the log-fold change of 8 transcripts with lowest adjusted p-value in the interaction analysis of FGF23 effect in diseased vs. FGF23 effect in healthy mice, 2×2 factorial design. N=5 (A), 10 (B) or 4 (C-H) biologically independent replicates per group.

Systemic proinflammatory effects of FGF23 have been reported to be mediated via the liver where FGF23 drives cytokine expression via FGFR4.^8^ However, we hypothesized that also renal tissue-resident cells could contribute to pro-inflammatory actions of FGF23. Therefore, we employed an isolated *ex vivo* model system and studied FGF23 signaling in precision-cut kidney slices (PCKS) (**Figure 6A**) from mice subjected to a classic model of adenine-induced nephropathy with advanced tubulointerstitial fibrosis (**Figure 6B**) and elevated concentrations of circulating intact and C-terminal FGF23 (**Supplemental Table 1**). In an RNAseq analysis of PCKS from these mice, a 24h treatment with FGF23 compared to vehicle resulted in 1 transcript with significantly increased expression, namely *Ccl4* encoding a proinflammatory macrophage/fibroblast chemokine (**Figure 6C, Supplemental Table 5**). GSEA of the Reactome database revealed no significant FGF23-induced gene set alterations in PCKS (**Figure 6D**), but GSEA by Wiki Pathways (WP) showed enrichment of Toll-like receptor signaling (**Figure 6E**) and Pathway Interaction Database (PID) showed enrichment of the Interleukin-12 pathway (**Figure 6F**), a pro-inflammatory early cytokine that leads to secondary release of interferon-gamma by natural killer and T cells.^37^ Overall, these data demonstrate evidence of kidney-intrinsic pro-inflammatory responses to FGF23 in an *ex vivo* PCKS model of diseased mouse kidney.

**Figure 6.**
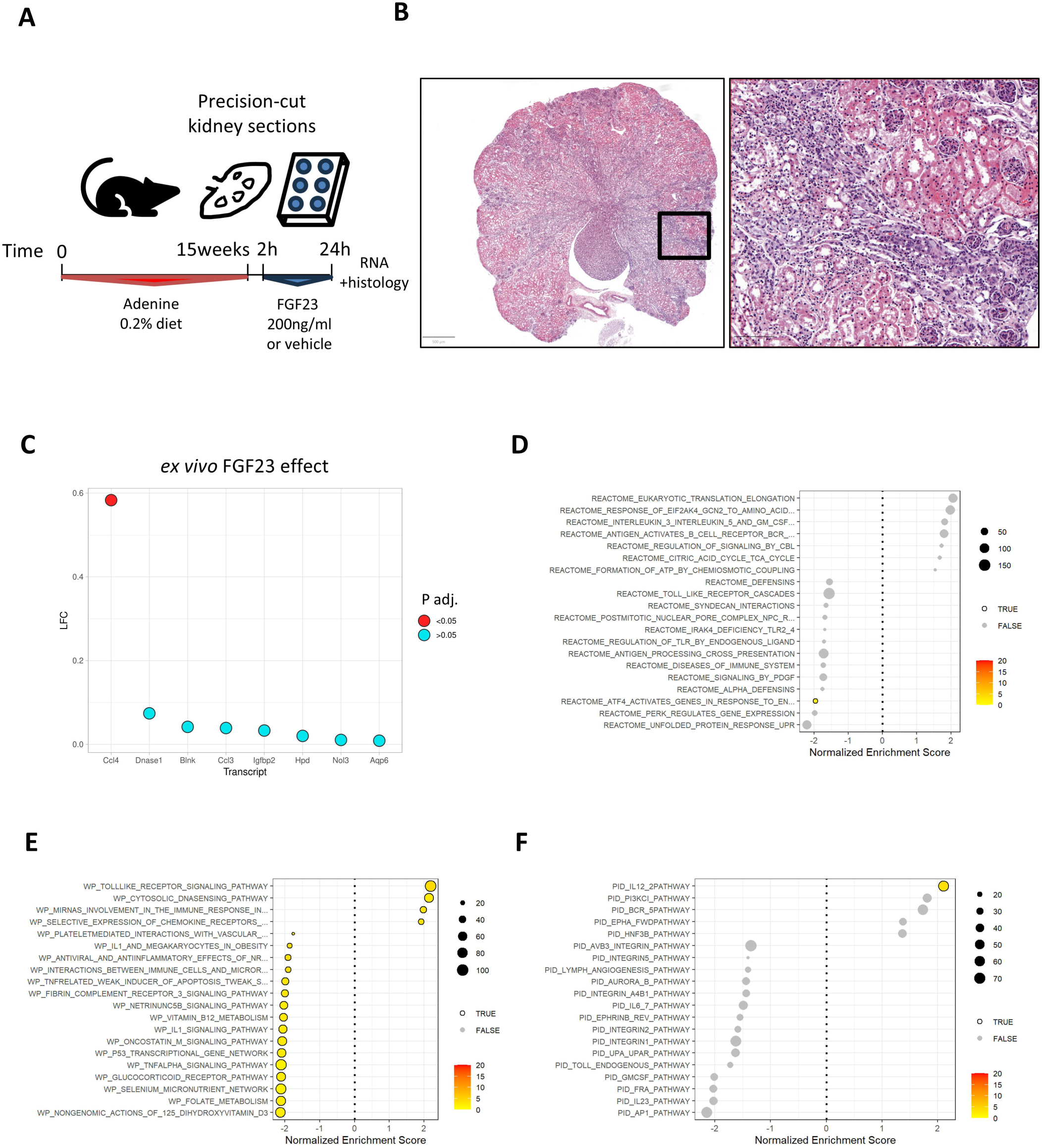
Intrarenal pro-inflammatory effects of FGF23 applied ex vivo in precision-cut kidney slices. Male DBA/2J mice underwent dietary treatment with 0.2% adenine for 15 weeks, followed by organ collection and 300µm precision-cut kidney slices (PCKS) for a 24h treatment with FGF23 or vehicle *ex vivo*. B depicts the fibrotic changes in a representative 4µm section of PCKS stained with hematoxylin-eosin (left scale bar, 500µm, right: 100µm). C shows the log-fold change of upregulated transcripts with lowest adjusted p value in FGF23 vs. vehicle comparison. D-F shows gene set enrichment analyses of FGF23 effects in Reactome (D), WikiPathways (E) and Pathway Interaction Database (PID) (F) gene sets. N=4 biologically independent replicates per group.

### Circulating FGF23 concentrations are associated with renal immune cell infiltration in human IgA nephropathy

Finally, we aimed to decipher if the link between elevated FGF23 and renal immune cell content found in mice can be translated to human kidney disease. To this end, we reassessed a sub-cohort of 59 patients with IgA nephropathy, a glomerular disease with varying degrees of renal inflammation and dysfunction whose kidney biopsies had undergone routine histopathological examination by board-certified pathologists, and transcriptome profiling of the non-glomerular (tubular) kidney fraction by RNAseq. Patient characteristics are described in **Supplemental Table 6**. We first reanalyzed and annotated a single-nucleus RNAseq dataset as reference (**Figure 7A, Supplemental Figure 5**) to infer renal immune cell content. As a validation step, we verified that transcriptome-inferred immune content of an immune cell/fibroblast cluster and of a macrophage cluster were associated with histopathological immune cell infiltration (**Supplemental Table 7**). We also verified the inverse association between FGF23 and renal function (**Figure 7B**). We then tested the association between transcriptome-inferred immune cell content and circulating FGF23. There were associations between both transcriptome-inferred immune cell content and FGF23, predominantly in those with moderate to advanced renal dysfunction (**Figure 7C-F**). In addition, we found associations between circulating FGF23 and several histopathological parameters of glomerular injury and tubulointerstitial immune infiltration in univariable or adjusted analyses (**Supplemental Table 8**). Overall, the present data provide evidence of a link between FGF23 excess and immune cell infiltration in IgA nephropathy.

**Figure 7.**
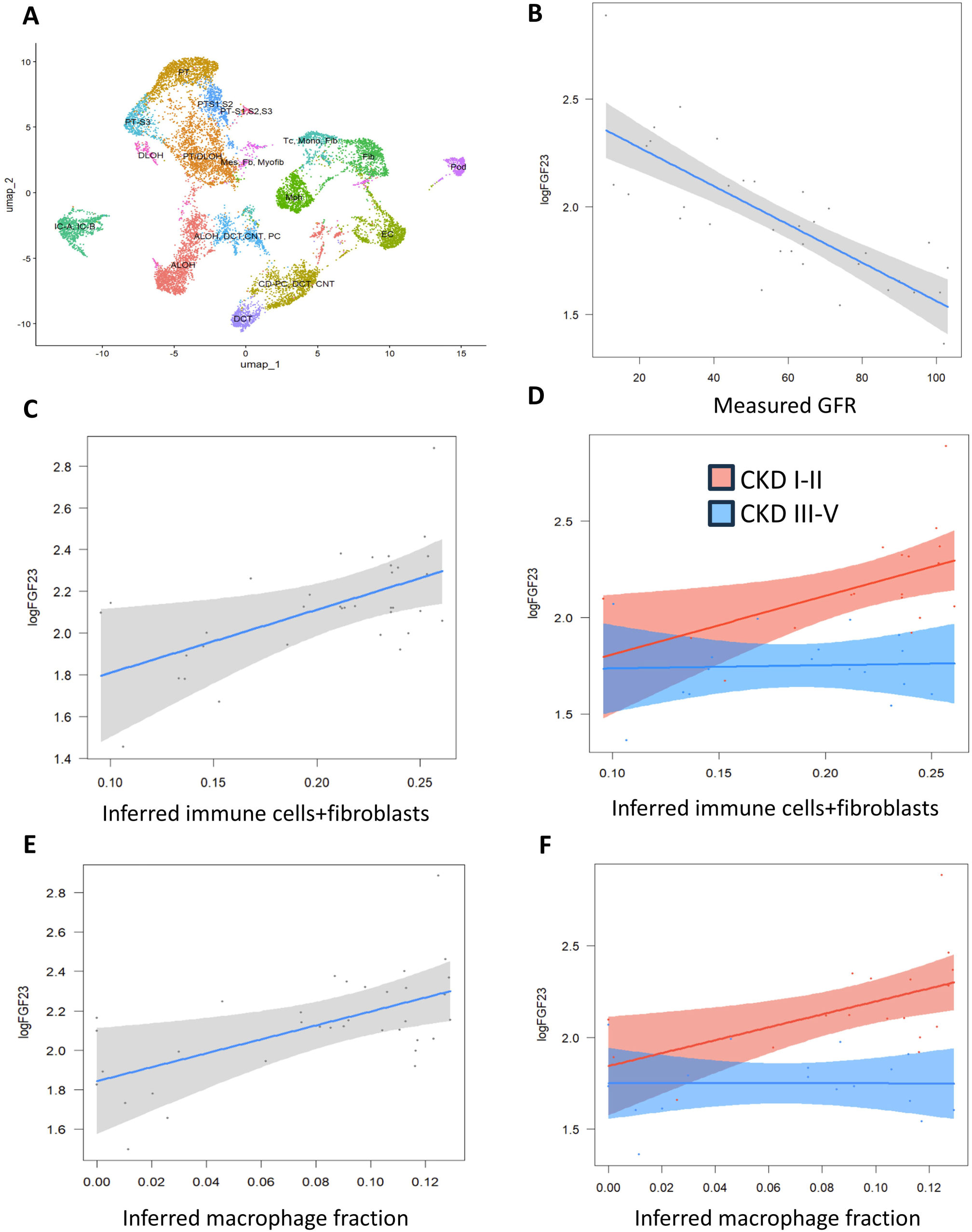
FGF23 associated with renal immune cell content in human patients with IgA nephropathy. As a reference, single-nucleus RNAseq data from 5 human kidney biopsies (GSE199711) of 2 healthy controls and 3 patients with chronic kidney disease (CKD) were reanalyzed and annotated (A), see also Supplementary Figure 5. 35 patients with IgA nephropathy from the Karolinska Kidney Biopsy Cohort showed an inverse univariable association between circulating FGF23 and measured glomerular filtration rate (GFR) (B). The associations between transcriptome-inferred renal fibroblasts and immune cells (C-D) or macrophages (E-F) and circulating FGF23 are shown, with adjustment for GFR, 25OH-Vitamin D and parathyroid hormone. D and F show disaggregated data stratified by GFR in CKD stages I-II and III-V.

## Discussion

In this study, we set out to describe the pathophysiological effects of FGF23 signaling kidney disease, with the intention to detect quantitative and qualitative aberrations in FGF23 signaling. In healthy mice, we detected no transcriptional alterations in the kidney at the 24h interval after the last treatment with FGF23 compared to vehicle. However, mice with Anti-GBM disease displayed proinflammatory cytokine expression and an excessive renal abundance and/or activity of macrophages and overall immune cells after the six-day regimen of FGF23 compared to vehicle. An in-depth transcriptomic analysis of ligand-receptor interactions in the kidney revealed FGF23-driven overactive collagen-integrin signaling that is associated with inflammation ^38^, and additional signaling involving mediators of tissue damage and acute kidney injury (AKI)-to-CKD transition such as Pdgfrb and Vcam1.^39^ Over long term, a genetically induced excess of FGF23 in two transgenic mouse strains increased renal macrophage content even in the absence of kidney disease. Finally, even *ex vivo* experiments in PCKS of kidney isolated from mice with adenine nephropathy revealed increased expression of *Ccl4* encoding a proinflammatory macrophage/fibroblast chemokine associated with cardiovascular events and disease,^40^ and transcriptional signatures of inflammatory signaling. These findings were preserved in associations between circulating FGF23 and renal immune cell content in human patients with advanced kidney disease.

Overall, the present data point towards a pathogenic potential of FGF23 excess over long-term duration or in kidney disease in a variety of settings including patients, three experimental mouse strains, different sexes and FGF23 administration routes and dosing (**Figure 8**). Singh et al. have previously reported that FGF23 induces FGFR4-mediated calcineurin signaling in the liver that then contributes to hepatic C-reactive protein expression and systemic inflammation.^8^ However, using the present approach of PCKS, our obtained results suggest that FGF23 may have direct kidney-specific adverse consequences in diseased kidney tissue, even in an experimental set-up of PCKS bypassing the proinflammatory systemic effects of FGF23 in the liver. Further, we previously reported rapid oxidative changes occurring in kidney tissue as a result of FGF23 administration,^22^ which may additionally promote a proinflammatory environment when occurring in kidney disease, a disease known to cause profound oxidative stress at the tissue level.^41^ Finally, our findings demonstrate a time dependency of FGF23 excess, with no significant global pro-inflammatory GSEA findings in short-term FGF23 treatment in mice with or without kidney disease. However, chronic FGF23 excess caused transcriptional evidence of renal macrophage overabundance even in mice without kidney disease. This correlates well with the pathophysiology observed in patients with AKI, where initial FGF23 excess was predictive of future transition to CKD.^42^

**Figure 8.**
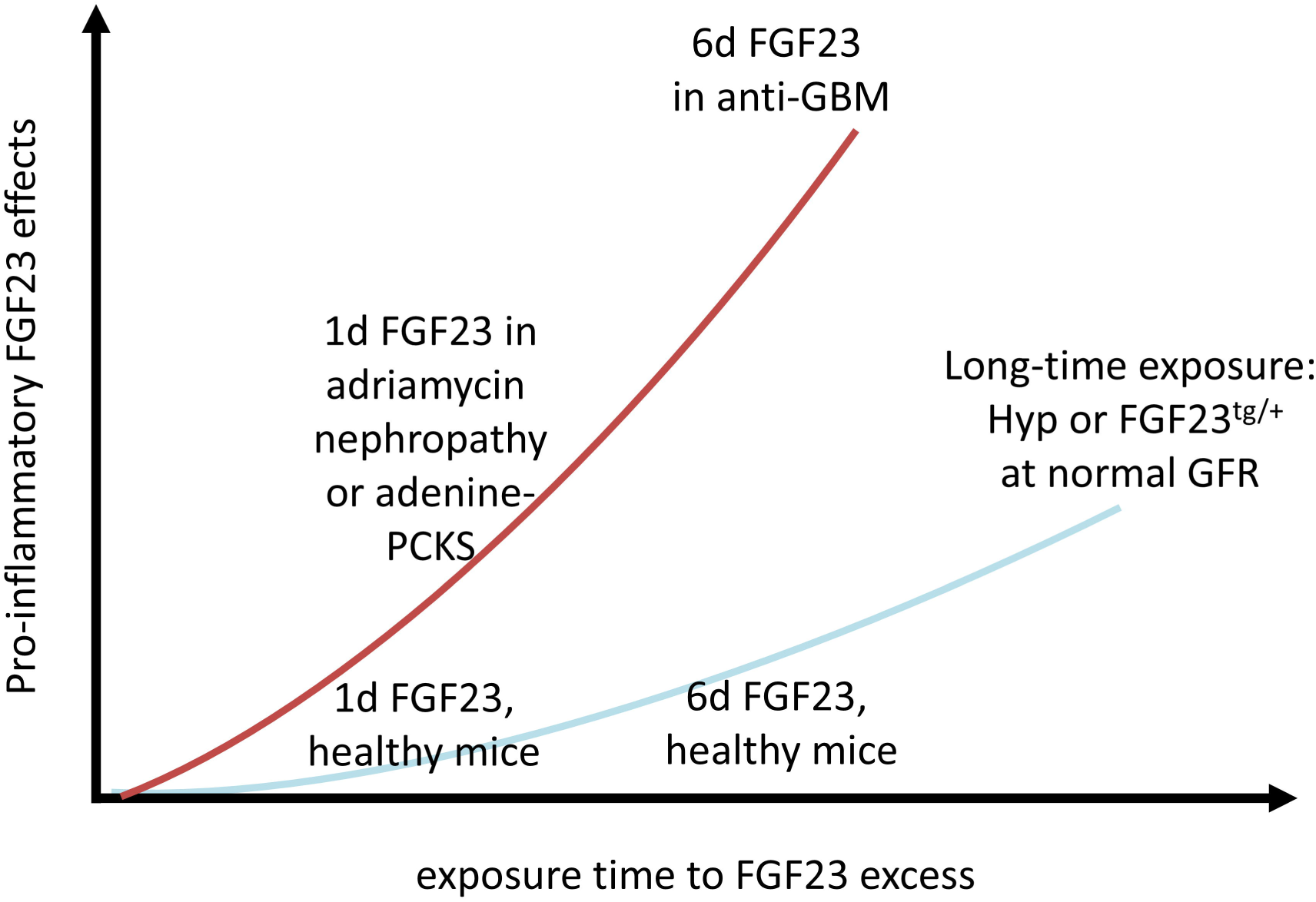
Model of FGF23 excess-driven renal inflammation. The graphical model integrates the obtained data and illustrates the tendency towards stronger pro-inflammatory effects in kidney disease models (red curve) and mice with normal kidney function (blue curve) that rises with increasing exposure time of FGF23 excess. Of note, human data are not included in this graph because causality of the observed associations cannot be determined.

A large body of evidence demonstrates that FGF23 excess is associated with adverse cardiovascular outcomes and mortality.^15,43,44^ Mechanistically, recent data by our group also point towards an adverse pro-inflammatory effect of FGF23 in experimental atherosclerosis.^10^ As a main implication of the present study, our findings would support the idea that it may be beneficial to reduce the amount of circulating FGF23 excess in kidney disease in order to prevent its toxicity. However, such a suggestion has to be cautiously approached, as FGF23 neutralization was shown to increase phosphate toxicity and ultimately mortality in rats with CKD.^13^ Further, recent data point towards potential benefits of a c-terminal FGF23 increase at the expense of intact FGF23 in experimental models, such as for uremic phenotypes of cardiomyopathy^6^ and anemia.^45^ Finally, one of the classical mainstays of CKD-MBD therapy is dietary phosphate restriction which ultimately protects from anemia and sarcopenia in experimental CKD,^7^ and metabolic acidosis and mortality in experimental AKI.^46^ In human patients with CKD, elevated serum phosphate is associated with an increased risk of death and cardiovascular disease.^47^ However, simply recommending patients with advanced CKD to reduce their phosphate intake did not improve mortality.^48^ Phosphate restriction induced by phosphate-binding agents is associated with lower overall mortality in in end-stage kidney disease,^49^ whereas in pre-dialysis populations, the effect of phosphate binders on patient outcomes remains so far inconclusive.^50^ As hyperphosphatemia is one of the key drivers of FGF23 excess in CKD, it is crucial to not only improve phosphate toxicity but also at the same time the presumed inflammatory FGF23 toxicity in order to ultimately improve clinical outcomes in CKD-MBD.

Strengths of the present study include the variations in animal strains, sexes and FGF23 application, in addition to the in-depth histological, transcriptomic and cytokine sub-proteome profiling in combination with advanced bioinformatics analyses. We chose relevant disease contexts in both murine and human data, and we cross-validated key findings by immunofluorescence and histopathology. However, this study also has some limitations. First, in wet-lab work we used human FGF23 in order to be able to directly quantify the injected recombinant protein by ELISA. This protein has its expected physiological effects in mice, and although human FGF23 is used in many mouse studies,^27,51–53^ human FGF23 might theoretically induce an allogenic immune response. However, an allogenic immune response would be expected to occur in all FGF23-treated experimental groups, which was not the case. Additionally, we validated our findings also in renal transcriptomes of Hyp mice that have excessive levels of endogenous murine FGF23.^28^ Next, we observed a discrepancy between transcriptome-inferred and immunofluorescence-estimated cell fractions. This may be in part attributable to the fact that our bulk deconvolution procedure was performed using healthy mouse reference single-cell RNAseq data, and therefore some adaptive or injured epithelial cells of the bulk transcriptomes could have been misclassified immune cells based on their transcriptional profile resembling rather immune cells. However, using standard markers CD45 and F4/80 in immunofluorescence, we confirmed the main trend of increased immune cells and macrophages induced by FGF23 excess in the Anti-GBM model. Further, the human dataset is of relatively small sample size and included a one-time measurement of FGF23 only. Finally, in the present study we almost exclusively focused on the kidney, neglecting the pro-inflammatory effects of the liver under FGF23 excess (SINGH), potential effects of phosphate excess on gut microbiota that can additionally generate bacterial-derived uremic toxins that may promote cardiovascular morbidity,^54^ and we did not engage with the direct renal tubular phosphate toxicity which FGF23 excess may promote by increasing phosphate concentrations renal tubular lumen.^44^

As a consequence of the obtained data, there is an incentive for future studies to investigate potential protective effects for uremic complications by carefully diminishing inflammatory FGF23 signaling both in CKD but also in the context of AKI to prevent the AKI-to-CKD transition.

To conclude, we employed several complementary approaches to determine adverse renal cell signaling induced by FGF23 in kidney disease, and we uncover substantial kidney-intrinsic pro-inflammatory effects induced by FGF23 and thereby further highlight FGF23 excess as a potential therapeutic target in CKD-MBD.

## Acknowledgements

The authors are thankful for the animal caretakers, staff of the FENO Morphological Phenotype Analysis Core for histology services, contributing staff and patients of the Karolinska Kidney Biopsy Cohort, and staff of the Bioinformatics and Expression Core Facility of Karolinska Institutet for sample processing and RNAseq.

## Funding

This work was funded by the Swiss National Science Foundation (grant no. 214187) and Karolinska Institutet Research Foundation (towards MBM). Work of MBM, HO and JP was funded by NovoNordisk Nordic Foundation. HO was funded by Westman foundation, Gelinstiftelsen, CIMED and Njurfonden. GGK was funded by Strategic Research Programme in Diabetes and Njurfonden.

## Conflicts of interest

Jaakko Patrakka’s research laboratory was financially supported by AstraZeneca and Guard Therapeutics International. Mikhail Burmakin was financially supported by Guard Therapeutics International. Annette Bruchfeld received consultancy fees from Amgen, AstraZeneca, Boehringer Ingelheim, CSL Vifor, Otsuka and Sobi and payment or honoraria for lectures, presentations, speaker’s bureaus, manuscript writing or educational events from AstraZeneca, Bayer, Boehringer Ingelheim, ChemoCentryx, CSL Vifor, Fresenius, GlaxoSmithKline and Otsuka. All other authors declare no competing interests.

## Accession codes

The murine bulk RNAseq datasets generated within this study are accessible at Gene Expression Omnibus under the accession number GSE272289. Human patients’ bulk RNAseq data have previously been deposited under the accession number GSE199711. Further datasets used in this study were GSE107585, GSE199711, GDS3361 and GDS879 as described in the methods section. Data from the Karolinska Kidney Biopsy Cohort can be requested from our Research Data Office at Karolinska Institutet.

## Notes

### Summary of Updates

Figure 7, labels of CKD stages I-II and III-V were corrected.

