## Supplemental Figures for "The renal response to FGF23 shifts from phosphaturia towards inflammation in kidney disease"

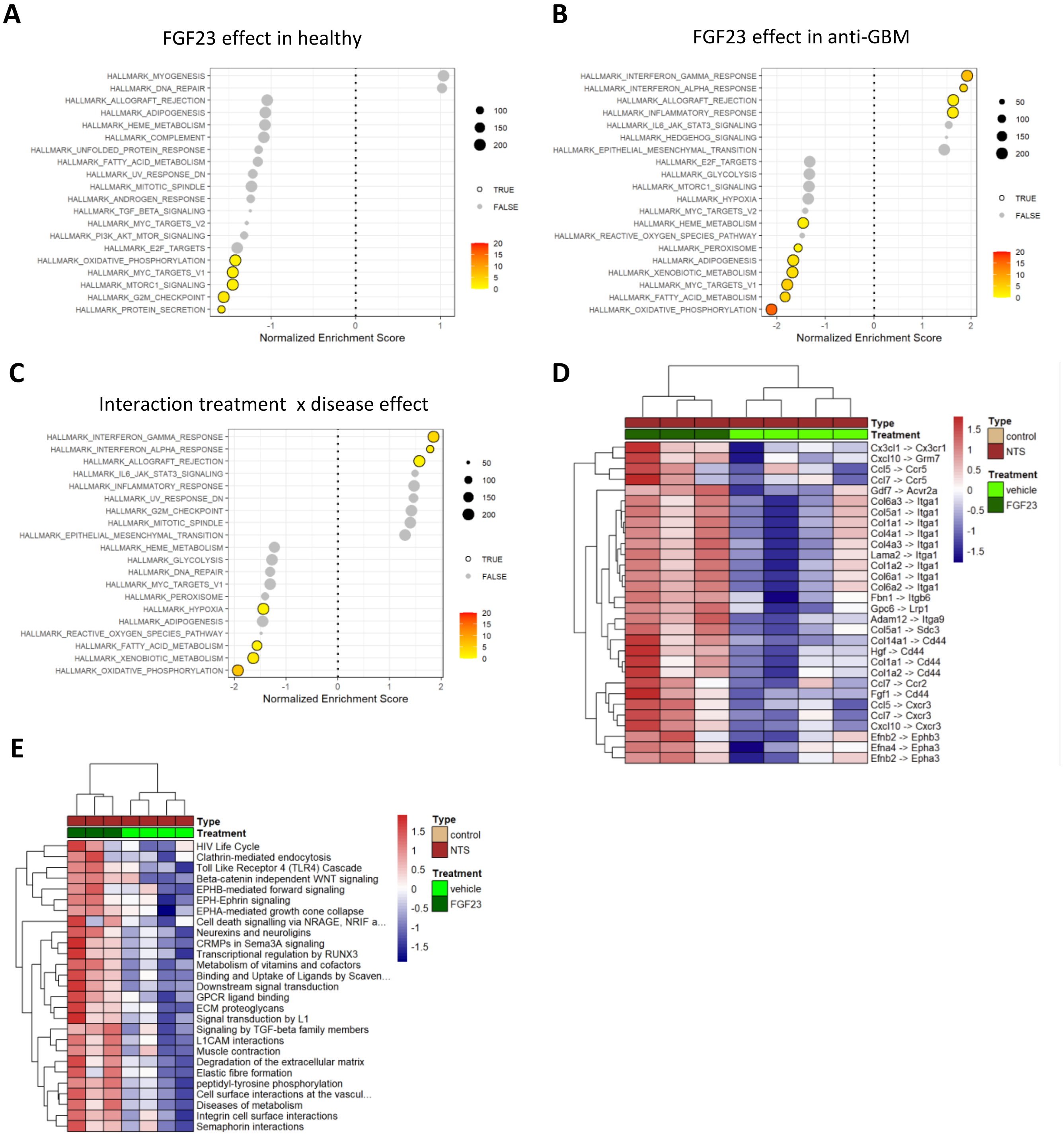


Supplemental Figure 1. Hallmark gene set enrichment analysis of FGF23 effects in healthy male C57BL/6 (A), mice with anti-glomerular basement membrane disease (anti-GBM, B), and interaction between FGF23 treatment and disease effect (C) using renal RNAseq. Significant ligand-receptor interaction pairs (D) and pathways (E) in renal bulk RNAseq of mice treated with nephrotoxic serum (NTS) to induce anti-GBM, followed by 6 days of daily recombinant FGF23 or vehicle injections.

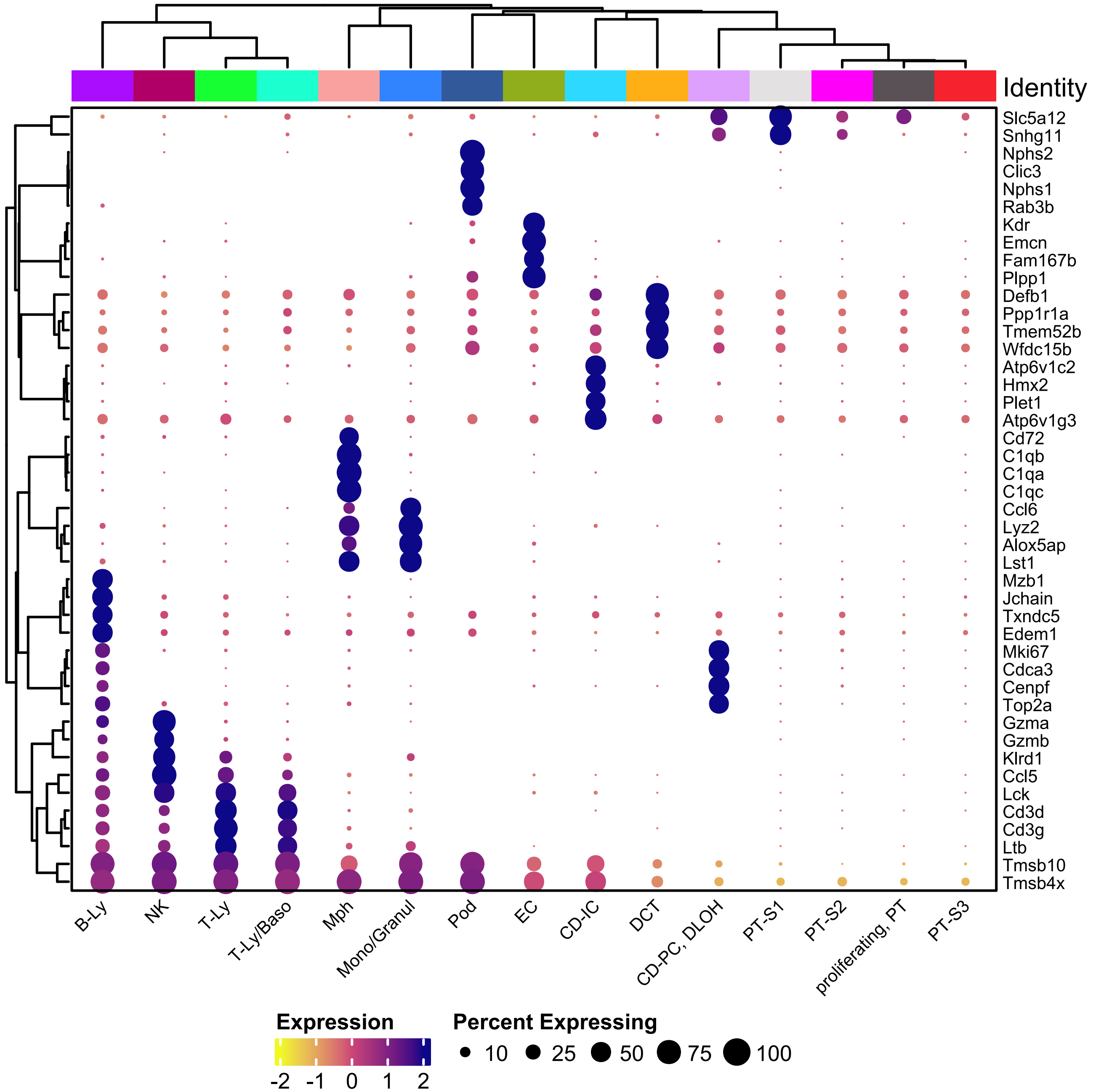


Supplemental Figure 2. Top four transcriptional markers clusters for each of 15 manually annotated clus- ters of kidney from 7 healthy sex-mixed C57BL/6 mice of single-cell RNAseq dataset GSE107585. Ly, lymphocytes. NK, natural killer cells. Baso, basophils. Mph, macrophages. Pod, podocytes. EC, endothelial cells. CD-IC, collecting duct – intercalated cells. DCT, distal convoluted tubule. PC, principal Cells. DLOH, descending loop of Henle. PT, proximal tubule. S, segment.

**A**

| | A | B | C | D | E | F | G | H | I | J | K | L |
|---|---|---|---|---|---|---|---|---|---|---|---|---|
| 1 | POSITIVE | POSITIVE | NEGATIVE | NEGATIVE | Blank | BLC | CD30L | Eotaxin | Eotaxin-2 | FAS ligand | Fractalkine | G-CSF |
| 2 | POSITIVE | POSITIVE | NEGATIVE | NEGATIVE | Blank | BLC | CD30L | Eotaxin | Eotaxin-2 | FAS ligand | Fractalkine | G-CSF |
| 3 | GM-CSF | IFN-gamma | IL1-alpha | IL1-beta | IL2 | IL3 | IL4 | IL6 | IL9 | IL10 | IL12-p40/p70 | IL12-p70 |
| 4 | GM-CSF | IFN-gamma | IL1-alpha | IL1-beta | IL2 | IL3 | IL4 | IL6 | IL9 | IL10 | IL12-p40/p70 | IL12-p70 |
| 5 | IL13 | IL17 | I-TAC | KC | Leptin | LIX | Lymphotactin | MCP-1 | M-CSF | MIG | MIP-1-alpha | MIP-1-gamma |
| 6 | IL13 | IL17 | I-TAC | KC | Leptin | LIX | Lymphotactin | MCP-1 | M-CSF | MIG | MIP-1-alpha | MIP-1-gamma |
| 9 | RANTES | SDF-1 | TCA-3 | TECK | TIMP-1 | TIMP-2 | TNF-alpha | sTNF RI | sTNF RII | Blank | Blank | POSITIVE |
| 10 | RANTES | SDF-1 | TCA-3 | TECK | TIMP-1 | TIMP-2 | TNF-alpha | sTNF RI | sTNF RII | Blank | Blank | POSITIVE |

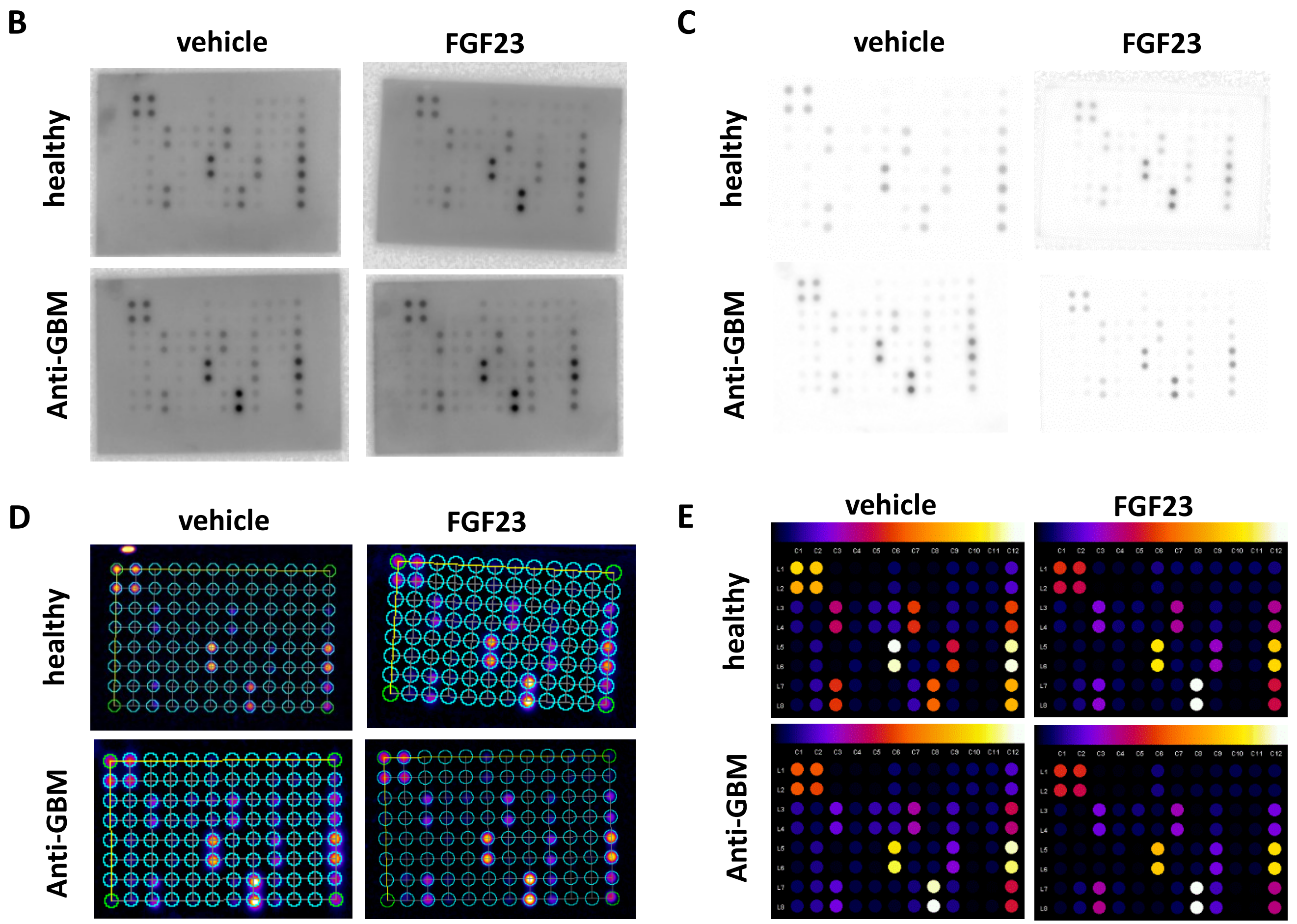


Supplemental Figure 3. Serum cytokine profiling using a colorimetric array for 40 murine proteins in doublets as indicated (A). One representative image per group is shown for digitally raw array scans (B), linear background subtraction (C), autosetting and semi-automated drawing of a grid (D), and quantification indicated by color scale with ascending values from left to right (E). N=4 biological replicates per group. Red rectangle in A shows the two proteins displayed in Figure 4E-F.

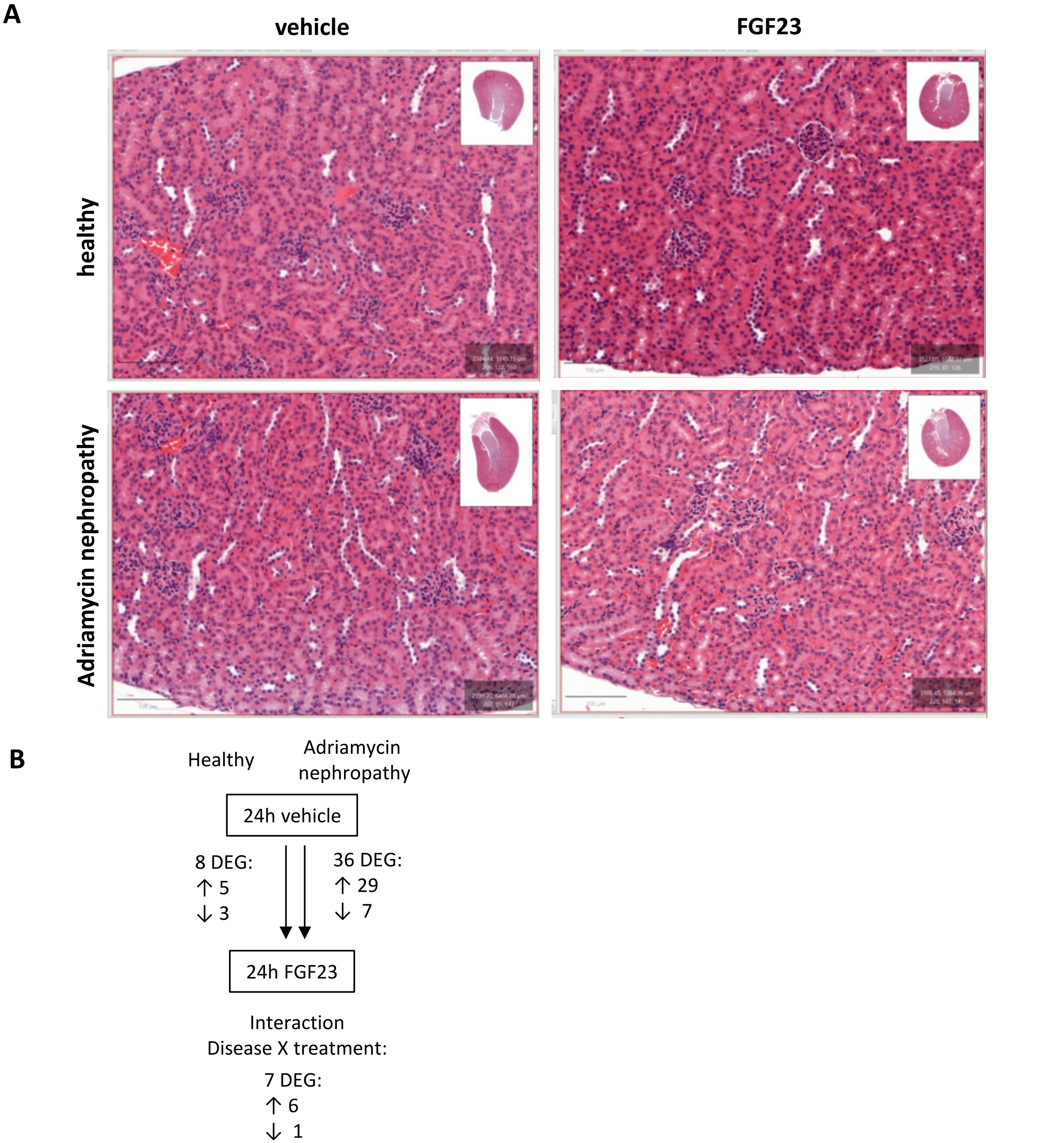


Supplemental Figure 4. Histological assessment of one representative kidney section per group, using Hematoxylin-eosin staining (A). N=4 per group of female BALB/c mice. B displays the number of differentially expressed genes (DEG) in renal bulk RNAseq comparing FGF23/vehicle treatment in healthy mice, in mice with Adriamycin nephropathy, and the interaction between disease state and FGF23/vehicle treatment.

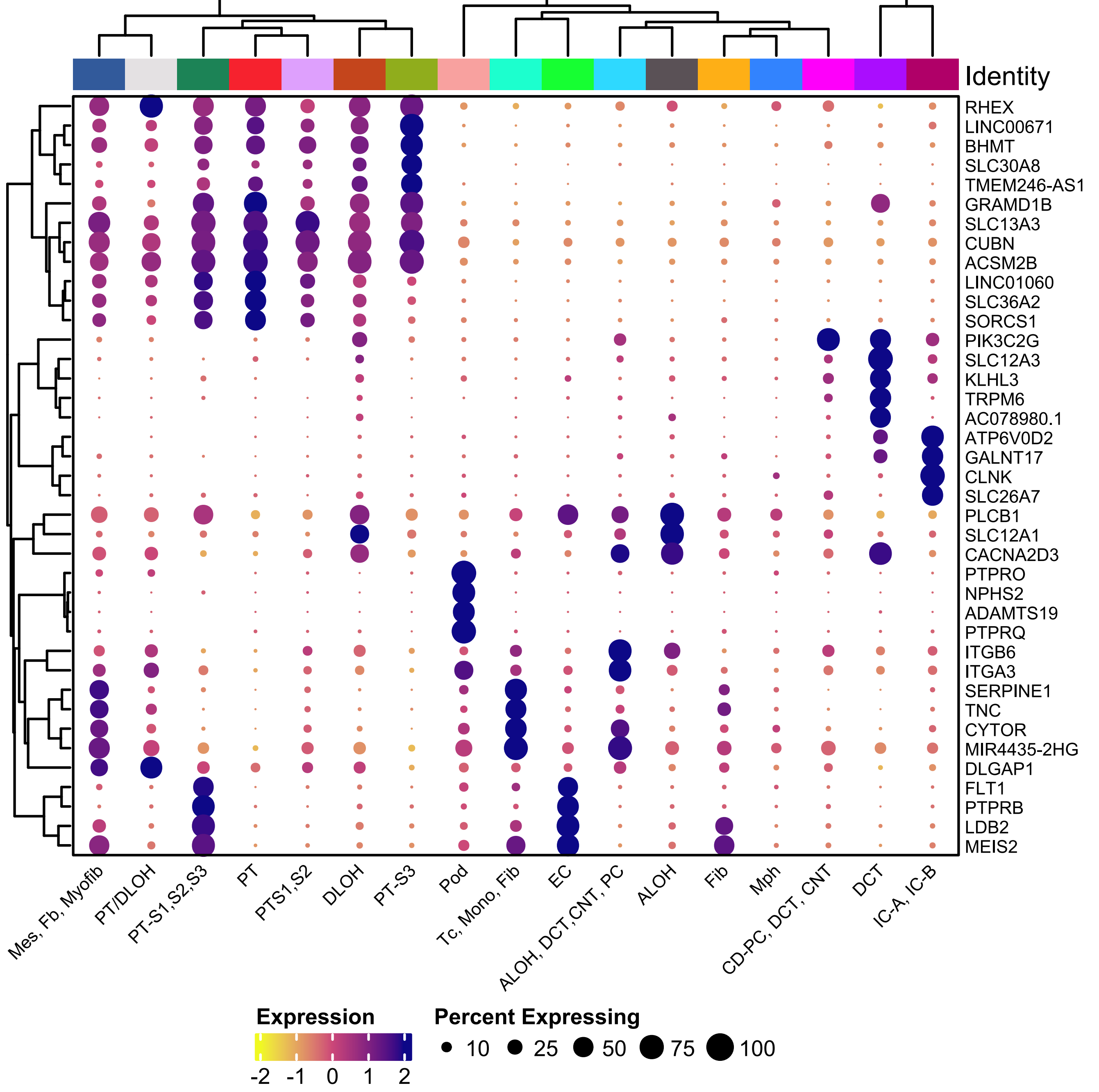


Supplemental Figure 5. Top four transcriptional markers for each of 17 manually annotated clusters of kidney single-nucleus RNAseq dataset GSE199711 from kidney biopsies of 3 patients with chronic kidney disease and 2 controls. Mes, mesangium, Fib, Fibroblast. Myofib, Myofibroblast. PT-S, Proximal Tubule Segment. DLOH, Descending Limb of Henle. Pod, Podocyte. Tc, T-cells. Mono, Monocytes. ALOH, Ascending Limb of Henle. DCT, Distal Convoluted Tubule. CNT, Connecting Tubule. PC, Principal Cells. Mph, Macrophages. CD, Collecting Duct. IC, Interalated Cells.
